# Crimean–Congo Haemorrhagic Fever Virus Circulates within Broad Ecological Networks of Ticks and Vertebrates

**DOI:** 10.1101/2025.11.20.689440

**Authors:** Agustín Estrada-Peña, Oliver Carnell, Sara R. Wijburg, Maya Holding, Hein Sprong

**Author notes:** Corresponding author (AEP). Current address: CSAI-Foundation, Ministry of Human Health, Madrid, Spain.

## Abstract

We produced spatial datasets of the known distribution of *Orthonairovirus haemorrhagiae* (formerly Crimean-Congo haemorrhagic fever virus, CCHFV) by compiling human cases, virus isolations, and serological data from humans and animals, spanning from Europe to southern Africa, with the aim on virus range modelling.

Models based solely on climatic variables produced unrealistic and overly extensive predictions, overestimating suitability in northern Europe and confirming that climate alone poorly explains CCHFV range. Approaches using only tick species distributions underperformed in many parts of Africa, reflecting the complexity of vector–host interactions and the absence of a strict tick–virus association.

The integration of human-biting tick distributions, livestock density, and chorotypes yielded the most robust results, accurately capturing over 90% of known occurrences, by identifying vertebrate assemblages most likely to amplify the virus while reducing dimensionality. Chorotypes, representing clusters of co-occurring hosts, enhanced model performance and improved the delineation of both the northern and African ranges limits of the virus.

Our results support a generalist epidemiological model in which multiple tick species, together with a broad range of vertebrate hosts, sustain CCHFV circulation. This ecological flexibility, rather than strict vector specificity, likely explains the CCHFV wide biogeographical range and occasional lineage expansions across continents.

This study provides the most comprehensive assessment to date of the abiotic and biotic determinants of the distribution of CCHFV. Although gaps remain, this study demonstrates that coupling abiotic and biotic predictors provides a more accurate and ecologically meaningful understanding of CCHFV distribution.

**Author Summary:** Crimean–Congo haemorrhagic fever virus (CCHFV) is a tick-borne pathogen causing severe disease in humans across Africa, Asia and Europe. Understanding where the virus may circulate is crucial for anticipating outbreaks and guiding surveillance. Previous studies have mostly relied on climate-based models, assuming that temperature and humidity determine where infected ticks can prevail. However, such models often overestimate risk, predicting virus presence in regions where no cases or vectors exist.

In this study, we combined climatic data with ecological information on ticks, livestock, and wildlife to create a more realistic prediction of CCHFV distribution. By incorporating host–vector interactions and vertebrate communities, our models accurately captured most known occurrences and revealed the ecological patterns underlying virus transmission. Our results suggest that CCHFV persists within broad networks of tick and vertebrate species rather than a single vector. This integrated approach offers a powerful framework to identify emerging risk areas and forms a good starting point to further improve our understanding of the ecological drivers of tickborne viral diseases.

## Introduction

Crimean-Congo hemorrhagic fever (CCHF) is a human disease caused by infection with *Orthonairovirus haemorrhagiae*, a segmented RNA virus commonly referred to as Crimean-Congo haemorrhagic fever virus, CCHFV, [1]. The virus circulates silently in tick-vertebrate-tick enzootic cycles with a brief viremia in vertebrates and prolonged viral maintenance in ticks by transstadial and transovarial transmission, and, less efficiently, by venereal transmission. The virus is widespread in the Old World, with a patchy distribution, and so is the reported disease occurrence, with the highest incidence in parts of Africa and West and Central Asia.

The main route of infection to humans is through the bite or handling of an infected tick of the family Ixodidae [2], in addition to infection through contact with infected livestock blood and tissues. Vertebrates that enable tick feeding, but do not support the multiplication of the virus, are known as maintenance hosts, as they contribute to the maintenance of the tick population.

Vertebrates in which the virus proliferate and infect ticks during feeding are known as amplification hosts, as they amplify the number of infected ticks in the environment. Cattle, goats, sheep and other large ruminants are effective amplification hosts and may infect ticks in the relatively short viraemic period [3,4]. Small mammals, like rodents and hares on which immature ticks feed, have a short and transient viremia, and are not considered very effective for a tick infection, but support the feeding of tick immatures. Co-feeding transmission of CCHFV has been demonstrated in the laboratory [5] but this way of amplification has been ignored in further field studies. Birds are most likely not viraemic, but some migratory birds may carry infected ticks in their trans-continental trips from Africa to Europe or Asia [6]. Studies support the limited role of long-distance migratory birds harbouring CCHFV-infected ticks for the establishment of the virus in a new territory [7]. Other routes of transmission involve the human to human (infected people admitted into hospitals with low protection for nursing personnel), from livestock carcasses via aerosol transmission during slaughter, or from crushing ticks manually.

Laboratory tests are the most straightforward method to assess the ability of ticks to acquire and transmit the virus. A review [8] found significant variability in results when using different virus strains, vertebrate hosts, or tick species. Despite these limitations and the limited number of tick species studied, these findings suggest that ticks from the genera *Hyalomma*, *Amblyomma*, or *Rhipicephalus* are vectors and reservoirs of the virus, capable of transmitting it trans-stadially and sometimes also trans-ovarially.

Laboratory vector competence studies enable the identification of ticks which can transmit the virus to hosts but cannot differentiate between primary and secondary vectors. Primary vectors of CCHFV are tick species which are necessary for maintenance of the circulation of the virus under natural conditions and transmit to humans. Secondary vector of CCHFV are tick species which can acquire and transmit the virus but cannot maintain virus circulations under natural conditions in the absence of a primary vector.

Since the last comprehensive review on CCHFV [6], our understanding of the virus epidemiology has seen minimal progress. CCHFV has been detected in 39 different tick species, collected from various hosts, while questing, or under experimental conditions, including *Amblyomma*, *Dermacentor*, *Hyalomma*, *Haemaphysalis*, *Rhipicephalus*, and even several species of soft ticks [9]. Interpreting these findings remains complex because ticks may acquire the virus from the blood of vertebrate hosts but may not be competent in transmission to other vertebrates or humans. For example, the role of soft ticks in the transmission has been demonstrated in the laboratory [10] and CCHFV RNA detected in the ticks, however this is not equivalent to transmission capacity [11]. Misconceptions about tick vectorial competence persist in studies aimed at evaluating the prevalence of the virus in ticks, humans, or animals [12].

Severe knowledge gaps in the ecology of CCHFV remain, regarding (a) which tick species maintain the circulation of CCHFV among amplification hosts, (b) which tick species infect humans, and (c) which vertebrates allow sufficient replication of CCHFV and feed ticks that could then be infected. To further complicate the matter, CCHFV can be genetically separated into six major clades or lineages: three predominantly found in Africa (lineages I–III), two in Europe (lineages V and VI), and one in Asia (lineage IV) [13]. Whether these lineages vary in their ability to infect different vertebrate or tick species (host range) or whether there are variations in the ability of ticks to transmit these different lineages is poorly understood but may have major impact on the current distribution of CCHFV and its ability to spread.

The production of maps pinpointing the range of CCHFV has been addressed on several occasions [14-18]. While it seems unlikely, it is not yet proven whether the range of CCHFV is exclusively linked to the known distribution of certain tick species or to a pattern of climate features without other explanatory forces. It is also unclear if the presence and/or abundance of specific wildlife species could represent the known range of CCHFV. A list of animals recorded to be positive in serological tests has been compiled [19], which was significantly updated [9]. Additionally, it is suggested that peculiarities in ecological relationships between ticks and vertebrates may influence the spatial complexity of the virus distribution [20]. A potential approach to understanding the CCHFV circulation and identifying areas prone to exposure could involve simultaneous mapping of vertebrates and ticks, along with climate and other relevant information.

We anticipate that utilising all available sources of information could produce suitable maps of the CCHFV range. Mapping the relationships between climate, ticks, and vertebrates across large regions as zoogeographical realms could facilitate the acquisition of epidemiological information through statistical interactions. This approach would not only benefit human health by predicting areas of risk for humans but also contribute to a deeper understanding of the actors involved in virus circulation. Such a framework should be grounded in extensive datasets of tick and vertebrate ranges, as well as sensitive methods that capture ecological associations and transform occurrence overlaps into an epidemiological message.

This study aims to model the distribution of CCHFV in the Western Palearctic and Tropics. It uses large georeferenced datasets of molecular and serological data from animals and humans, known georeferenced clinical records of the virus, and the joint distribution of 82 tick species and 121 genera of vertebrates reported as either maintenance or amplification hosts. The study focuses on detecting and defining the hypothetical environmental niche for the virus. If this environmental niche exists, CCHFV could only occur within the niche’s spatial projection, regardless of the presence of ticks and vertebrates in the area. The study also aims to identify the epidemiological associations of the virus with primary and secondary vectors. Additionally, it explicitly tests whether CCHFV occurs in conjunction with biogeographical constructs that describe ranges of cooccurring ticks and vertebrates as biogeographical entities known as chorotypes (Estrada-Peña et al., 2025). The data presented in this study are intended to support a comprehensive approach to simultaneously test the interactions of communities of ticks and hosts, climate, and livestock distribution to explain the geographic distribution of CCHFV.

## Methods

### Area of study

The target territory includes the Western Palearctic and the Tropics, between the coordinates 72°N, 36°S, 58°E, 18°W. The territories eastern to the Palearctic domain have not been included because the lack of adequate reporting of ticks, virus detection and/or human cases. Findings of CCHFV in the area are commonly referenced to large administrative divisions (e.g. provinces, without coordinates) therefore preventing modelling. This study involves the range of both ticks and vertebrates as explanatory variables for modelling CCHFV extent, that needs coordinates.

### Source of records and climate data

We consulted multiple sources to compile a reliable dataset of tick and vertebrate occurrences within the target territory. Tick records were sourced from three previous compilations [21, 22], available data for specific tick species in Europe [23-25], and GBIF [26]. The list of ticks included in this study is available as Supplementary Material 1. For vertebrate distributions, we utilized two previously compiled datasets [21, 22] to identify vertebrate species reported as tick hosts over the past 40 years. The list of vertebrates serving as tick hosts in the target territory was updated with published data [27] and additional data downloaded from GBIF [26].

Initially, the dataset included 132 tick species and 403 host species. Data on *Haemaphysalis* spp. were removed after initial tests demonstrating the lack of correlation of these species with records of CCHFV, parasitism to humans or solid links of ticks in the genus and transmission of the virus. To address the logistic challenges posed by modelling over 400 species of vertebrates, we downgraded the original dataset to genera of vertebrates instead of species. The dataset of ticks was examined to eliminate species with less than 50 records. In total, we analyzed a dataset containing 92,404 geo-referenced records of 82 tick species and 4,920,908 geo-referenced records of vertebrates across 121 genera.

### Climate data

Climate data were used to model environmental suitability for several groups of organisms across the target territory at a spatial resolution of 4 km / pixel. We sourced monthly data on maximum and minimum temperature and water vapor deficit from the TerraClimate repository (https://www.climatologylab.org/terraclimate.html, [28]) for the period between 1990 and 2020. For each month, we calculated the average values across the target territory using monthly values (1990–2020) and the “terra” package (v 1.8.70, [29]) for the R programming environment.

We performed harmonic regression on the 12 averaged monthly values (temperature or water vapor deficit) against time (in months, from 1 to 12). The coefficients derived from the harmonic regression were used as the explanatory variables for tick or vertebrate occurrence modelling [30]. These variables are orthogonal and thus non-self-correlated, eliminating the need to assess collinearity among variables for good modelling practices. They also retain ecological relevance for environmental suitability modelling and capture critical conditions for organism survival better than sets of pre-tailored climate data [31]. Nine variables were used to describe climate, namely the first three coefficients of the harmonic regression for maximum temperature (tmax), minimum temperature (tmin), and water vapor deficit (vpd).

### Modelling the distribution of ticks and vertebrates and building of chorotypes

We generated maps of the expected distributions of ticks and vertebrates, using the approach known as stacked species distribution modelling (SSDM [32]). This method addresses the simultaneous evaluation of multiple algorithms on the complete set of species (either ticks or vertebrates), selecting the best combination through ensemble modelling (i.e., combining the results of various algorithms). The range of each organism was modelled using the known records, thinned to the 4 km resolution of the explanatory variables. We employed the following algorithms: generalized additive model (GAM), multivariate adaptive regression splines (MARS), and support vector machines (SVM) in the SSDM package (v. 0.2.11 [32]) in R. We also used the implementation of the algorithm MaxEnt in the package “Wallace” (v. 2.1.3 [34]).

For GAM, we ran 500 iterations per species/genus with an epsilon value of 1E-8; for MARS, we allowed up to 4 degrees of interactions; for SVM, we used the same epsilon as for GAM, with three cross-validation steps. For MaxEnt we optimized the parameters of feature classes and the regularization parameters, as recommended [34, 35]. To select the best model parameters the feature class were restricted to lineal, quadratic, and product. The regularization multiplier values were 1, 2, and 5 [36]. Parameters not specified were kept at their default values in the package(s). In all cases, 70% of the records were used for model training and 30% for model testing. Pseudo-absences were generated automatically by the software in a number proportional (1:1) to the actual records of the organism to be modelled.

The metrics for selecting the best models for inclusion in the ensemble SDM were the threshold value (a balance between specificity and sensitivity), the omission rate (proportion of missing records with the best model), sensitivity (the proportion of correctly predicted presences), specificity (the proportion of correctly predicted absences), proportion of correctly detected records (overall accuracy), and the Cohen’s kappa. The area under the curve (AUC) was used for model comparison, although its interest as a metric for model performance was restricted to comparisons within the same species. The complete set of maps for both ticks and hosts is available in FigShare at https://figshare.com/s/9ffe21744531bddf67db.

The range maps of ticks and/or vertebrates were used to (1) establish relationships between the records of CCHFV and species of vectors and hosts and (2) calculate chorotypes as published by Real et al., 2008. A chorotype is a set of species that co-occur in a significant way resulting in a common range. Chorotypes are the roots of an epidemiological regionalization because they consist of the identification of spatial units with similar species composition [37]. Their applicability for exploring of epidemiological properties of co-occurring ticks and hosts in large territories was demonstrated before [38].

We applied a clustering procedure [39] based on Principal Components Analysis (PCA). Chorotypes resulted from optimizing the number of clusters required to explain observed variability through the ‘elbow’ method in the package “recluster” (v 2.9, [40]). The aim of PCA was both to reduce the spatial variability into axes that explained variance and to define epidemiological spatial units (chorotypes) that will be used later as part of the set of explanatory variables used to capture the range of CCHFV.

### Compilation of human cases

Data on the geographical distribution of CCHFV were used to train the models predicting the distribution (probability of presence) of CCHFV. The reported cases of CCHFV in the target territory were obtained from compilations [15, 16] that were complemented with known cases from Portugal, Spain, and Türkiye, not included in the mentioned compilations but reported previously [41] and with a few records available in Genbank with details to a locality of isolation. Most records of CCHFV in Genbank refer to large administrative divisions, commonly countries. The data reported to whole countries cannot be unequivocally ascribed to a pair of coordinates. The total number of georeferenced human cases and virus isolation was 725.

### Compilation of serological data

We compiled published data on serological surveys conducted in humans and animals by performing a comprehensive review of electronic bibliographic databases for English-language journals up to September 12^th^, 2024. The serological survey data served as an additional source for training the models of CCHFV distribution and are available in Supplementary Material 2.

The compilation itself includes papers that may lack suitable spatial details for mapping. For this paper, only studies with appropriate spatial information were included (see below for spatial processing of information). For papers containing suitable data, we recorded the following information: DOI, first author, title, year of publication, country, region of interest (e.g., a province, region, etc.), and the Global Administrative Areas (GADM) level 3 administrative division. GADM level 3 is equivalent to the municipalities (NUTS3) in Europe and is available at https://gadm.org.

We also included the year the study was conducted, the number of people (or animals) involved in the serological study, their occupation, whether tick bites were recorded, and whether they had contact with animals. Additionally, we compiled data on seroprevalence, specific IgG and IgM values, the test used for the serological survey, and the confirmatory test for the virus. If sequencing was performed, we included the GenBank accession number of the sequence(s). We included details of environmental factors, presumed risk factors, and how these factors were statistically evaluated. In the case of animals, we also included details about collections of feeding ticks and if the virus was detected in these ticks. Notably, most papers did not include all the attributes mentioned above. We did not remove papers missing one or several components to ensure a complete dataset of serological surveys conducted for CCHFV.

The inclusion of serological data for either animals or humans posed a challenge in terms of homogenisation. The literature search revealed a diverse range of formats. Serological data was most reported as data for administrative divisions of varying sizes, always smaller than the country, but rarely data was presented with spatial coordinates or the name of a locality that could be accurately referenced to a pair of coordinates. To address this issue, we simplified the original dataset of serological data to a common resolution, focusing on the smallest administrative divisions. Subsequently, we employed a tessellation method, which has proven effective in similar problems [22]. This method involved covering the entire target territory with hexagons, each with a radius of 2°. Each hexagon was then overlaid with the serological rates for either animals or humans using GIS software’s overlap routines. The values of serology, after being reported and transformed to the smallest administrative division, were assigned to the corresponding hexagon. The centroid of each hexagon within the honeycomb, where positive serological rates were detected (either for humans or animals) or where molecular tools indicated the presence of the virus, was considered the geo-referenced point for further modelling.

Data about human cases, virus isolations obtained from Genbank, and the range of serological surveys in either humans or animals, have been included in Figure 1.

**Figure 1.**
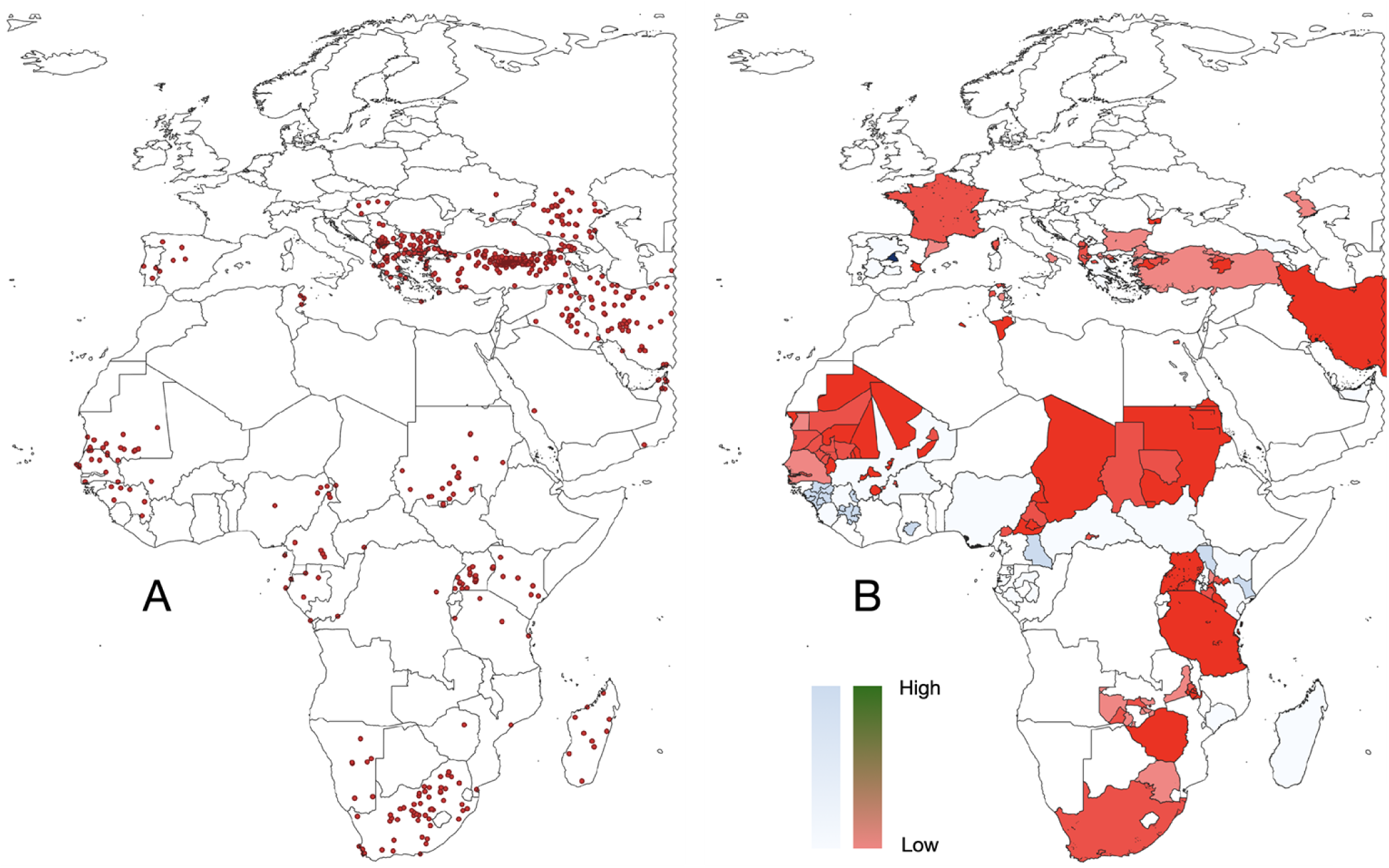
A. The 725 occurrences of Crimean-Congo haemorrhagic virus (CCHFV) detected in humans. B. The geographical distribution of the serological studies compiled from the literature, carried out in animals (tones of blue, 144 papers) or in humans (tones of red, 107 papers). The tones of either blue or red are representations of the serological prevalence.

## Modelling the range of CCHFV

### 1. The climate niche

We first determined if CCHFV has a climate niche. This was done by analyzing the 725 human case records and the 1,217 pseudo-absences records randomly placed over the target territory, against the nine environmental variables explained above. Climate information for each presence/pseudo-absence point was extracted. Next, we subjected the set of points to a Non Metric Multidimensional Scaling (NMDS). NMDS is a gradient analysis approach that uses ordination [42]. It compares the pairwise dissimilarity between the climate covariates of viral records using their variation among sites. Each record of CCHFV is assigned a pair of “coordinates” resulting from the NMDS, which provides it with a “position” in the classification space. The set of data is then subjected to a cluster analysis using kmeans. If the cluster analysis can statistically (a) separate negative and positive records, and (b) group all the presence points into a well-defined cluster, it is indicative of a climate niche for CCHFV because presences and pseudo-absences can be separated with an assumable error by the values of the environmental variables. Conversely, if the cluster analysis cannot resolve a single group of records, it is suggestive of the existence of multiple climate regions where CCHFV can thrive; in the later, it would be not recommendable to map the probability of distribution of CCGFV using only climate variables.

### 2. The tick niche

We modelled the distribution of CCHFV using only the tick distribution, without any prior assumptions about their vectorial capabilities. This approach suggests that any tick species could be involved in CCHFV circulation, and ranges of several species may overlap within the known CCHFV range. We assumed multiple tick species might be positively correlated with the known range, so we ranked their importance to create a preliminary rank of each tick in the model CCHFV-ticks. Additionally, we considered that some ticks might not actually be involved in CCHFV circulation but could overlap its known range, which we deemed as spurious correlations.

It is impractical to model the extent of CCHFV using raster layers representing the hypothetical involvement of 82 species of ticks, entered one by one, in pairs, triads, and so on (this represents the factorial of 82 species in terms of possible combinations). We employed a Random Forest (RF) framework to evaluate the relative contribution of each tick to the potential distribution of the virus. RF is an ensemble learning method based on the construction of many decision trees, each trained on a bootstrap sample of the original dataset [43]. Predictions from individual trees are aggregated to produce robust and stable estimates, reducing overfitting and capturing non-linear relationships between predictors (the tick ranges) and the response variable (presence/absence of CCHFV).

We trained the RF using the observed presence and pseudo-absence of viral cases as the binary response (725 points of CCHFV presence, and the set of 1,217 pseudo-absences) and vector species ranges as predictor variables. Each tree in the forest was grown without pruning, and a subset of predictors was randomly sampled at each split to decorrelate trees and improve generalization. Variable importance was assessed using permutation-based measures, which quantify the decrease in predictive performance when a given predictor is randomly permuted. The RF model outputs include the predicted probabilities of CCHFV occurrence, and the variable importance identify the most influential tick species.

We calculated the True Skill Statistic (TSS) that balances both omission (false negatives) and commission (false positives) errors. A high TSS indicates that the model accurately identifies both presences and absences. It ranges between -1 and 1, where 1 is the perfect agreement between predictions and observations, and <0 is worse than random. The Root Mean Squared Error (RMSE) measures the average magnitude of prediction errors. Lower RMSE means the predicted probabilities are closer to the observed outcomes. We calculated the LogLoss (Logarithmic Loss / Cross-Entropy Loss) that penalizes confident but incorrect predictions more heavily than small errors. It is particularly sensitive to probabilities near 0 or 1 that are wrong. In the context of ecological modelling, a low LogLoss indicates that the model not only classifies correctly but also assigns accurate probabilities of presence, which is critical when making spatial risk maps. Values near zero indicate perfect predictions. The model was used to map the expected distribution of CCHFV in the target territory using the raster layers of the species of ticks detected by the trained model as significant for the output.

### 3. Other models of CCHFV occurrence

We assumed that the previous modelling protocols may be simplistic representations of the range of CCHFV and aimed to explore various combinations of other explanatory variables. In each instance described below, we employed the algorithms GAM, MARS, SVM, and MaxEnt, with the same parameter tuning as before. The metrics used to select the best models for inclusion in the ensemble SDM included the threshold value (balance between specificity and sensitivity), the omission rate (proportion of missing records with the best model), sensitivity (proportion of correctly predicted presences), specificity (proportion of correctly predicted absences), overall accuracy (proportion of correctly detected records), and Cohen’s Kappa. In the case of the chorotypes as descriptive variables, we assumed that they encapsulated the redundant information contained in the raster ranges of several species, thereby decreasing the number of variables entered. Here, our hypothesis was that if a chorotype is positively associated with the occurrence of CCHFV, then some ticks/hosts in the chorotype may be involved in the transmission or amplification of CCHFV.

All combinations of variables included the nine environmental variables, and:

*1)* <u>The chorotypes of ticks and vertebrates, included separately as independent sets of variables</u>. This approach was intended to capture the joint effect of a variety of ticks and hosts as explanatory variables of the CCHFV occurrence. No single organisms were separately introduced in the model. The aim was to demonstrate that CCHFV occurs under given combinations of hosts and/or ticks.
*2)* <u>The distribution of human-biting tick species, the distribution of chorotypes of vertebrates, and the density of</u> <u>livestock</u>. This approach was intended to evaluate the distribution of human biting ticks, the distribution of hosts (as their chorotypes), and the density of livestock as explanatory variables. The rationale was that most of the transmission events of CCHFV are derived from tick bites to humans; some cases are known to be produced by aerosol transmission of the virus during peridomiciliary slaughtering or bushmeat practices.

The ticks included in this analysis were *Amblyomma hebraeum, Amblyomma variegatum, Dermacentor marginatus, Dermacentor reticulatus, Hyalomma aegyptium, Hyalomma anatolicum, Hyalomma excavatum, Hyalomma marginatum, Hyalomma rufipes, Hyalomma scupense, Hyalomma truncatum, Hyalomma turanicum, Ixodes ricinus*. Among these ticks, some are known to transmit CCHFV in the laboratory [8], while others have been repeatedly reported as parasites of humans [44] but have not been explicitly tested as vectors of CCHFV. The *Rhipicephalus sanguineus* group, known to bite humans, was excluded from this analysis because the probable misidentification of many specimens in the large range could bias the information about species that bite humans against those that do not. The information about livestock density was obtained from the world map of livestock provided by the Food and Agriculture Organization (FAO) (https://www.fao.org/livestock-systems/global-distributions/en/) and entered as three separate layers for cattle, sheep, and goats.

## Results

### Results on modelling of ticks and vertebrates

The performance metrics of the algorithms used to model the distribution of ticks and vertebrates are included in Table 1. Values represent the averages across the full dataset of either ticks or vertebrates, and do not refer to a particular species. Overall, model performance was consistently higher for ticks than for vertebrates, despite the substantially larger number of vertebrate records relative to those of individual tick species.

**Table 1.** The performance indexes for each algorithm, averaged across the complete set of tick species and hosts genera. These parameters include the threshold (optimised for sensitivity and specificity to produce a suitable habitat threshold), AUC (Area Under the Curve), the omission rate (proportion of presences classified as absences), sensitivity (model’s ability to detect positive instances), specificity (proportion of true negatives correctly identified), accuracy (percentage of correctly classified records, either presence or pseudoabsence), and Cohen’s Kappa (agreement between observations and predictions).

| Algorithm | Threshold | AUC | Omission rate | Sensitivity | Specificity | Accuracy |
| --- | --- | --- | --- | --- | --- | --- |
| Ticks |  |  |  |  |  |  |
| MAXENT | 0.2805 | 0.9578 | 0.4677 | 0.9012 | 0.5298 | 0.5322 |
| GAM | 0.5974 | 0.9639 | 0.0866 | 0.9136 | 0.9130 | 0.9133 |
| MARS | 0.6305 | 0.9322 | 0.0981 | 0.9034 | 0.9049 | 0.9082 |
| SVM | 0.7736 | 0.9476 | 0.1008 | 0.9001 | 0.8982 | 0.8991 |
| Vertebrates |  |  |  |  |  |  |
| MAXENT | 0.4360 | 0.8623 | 0.2151 | 0.7853 | 0.7847 | 0.7849 |
| GAM | 0.3970 | 0.8758 | 0.2039 | 0.7954 | 0.7967 | 0.7961 |
| MARS | 0.4350 | 0.8492 | 0.2152 | 0.7853 | 0.7843 | 0.7848 |
| SVM | 0.4207 | 0.9467 | 0.4249 | 0.8652 | 0.5699 | 0.5750 |

For ticks, the modelling generated predictive maps for 82 species, with generally strong performance (mean sensitivity = 0.91; mean specificity = 0.91). Generalised Additive Models and MARS achieved the highest accuracy, whereas SVM performed slightly less effectively. It is notable that the models developed using Maxent showed a poor balance between omission rate, specificity, and accuracy, as evidenced by their high omission rate. For vertebrates, the modelling of the known distributions yielded maps for 121 genera, with more modest results (mean sensitivity = 0.73; mean specificity = 0.73). Among the algorithms, MARS, GAM, and SVM performed best than MaxEnt, that outputted high omission rates. Notably, the lowest proportion of correctly allocated records of vertebrates occurred with SVM. The final output of the maps of each species/genus was prepared with the best combination of algorithms and variables.

### Chorotypes reduced the variance of the range of multiple species

Chorotypes are synthetic expressions of the co-occurrence patterns of either ticks or hosts, that explain the range of several taxa with similar distributions, detecting even clusters of organisms that co-occur "inside" other chorotypes. The original 82 distribution maps of ticks were reduced to four significant chorotypes (Figure 2), jointly explaining 81.24% of the total variance (49.26%, 14.65%, 10.03%, and 7.37%, respectively). The ordination of the genera of hosts along the first PCA-derived axes is shown in Supplementary Figure 2. The original 121 maps of hosts were reduced to three chorotypes (Figure 3) that explained 74.22% of the variance (47.24%, 20.86%, and 6.12%, respectively). The ordination of the genera of hosts along the first PCA-derived axes is shown in Supplementary Figure 2.

**Figure 2.**
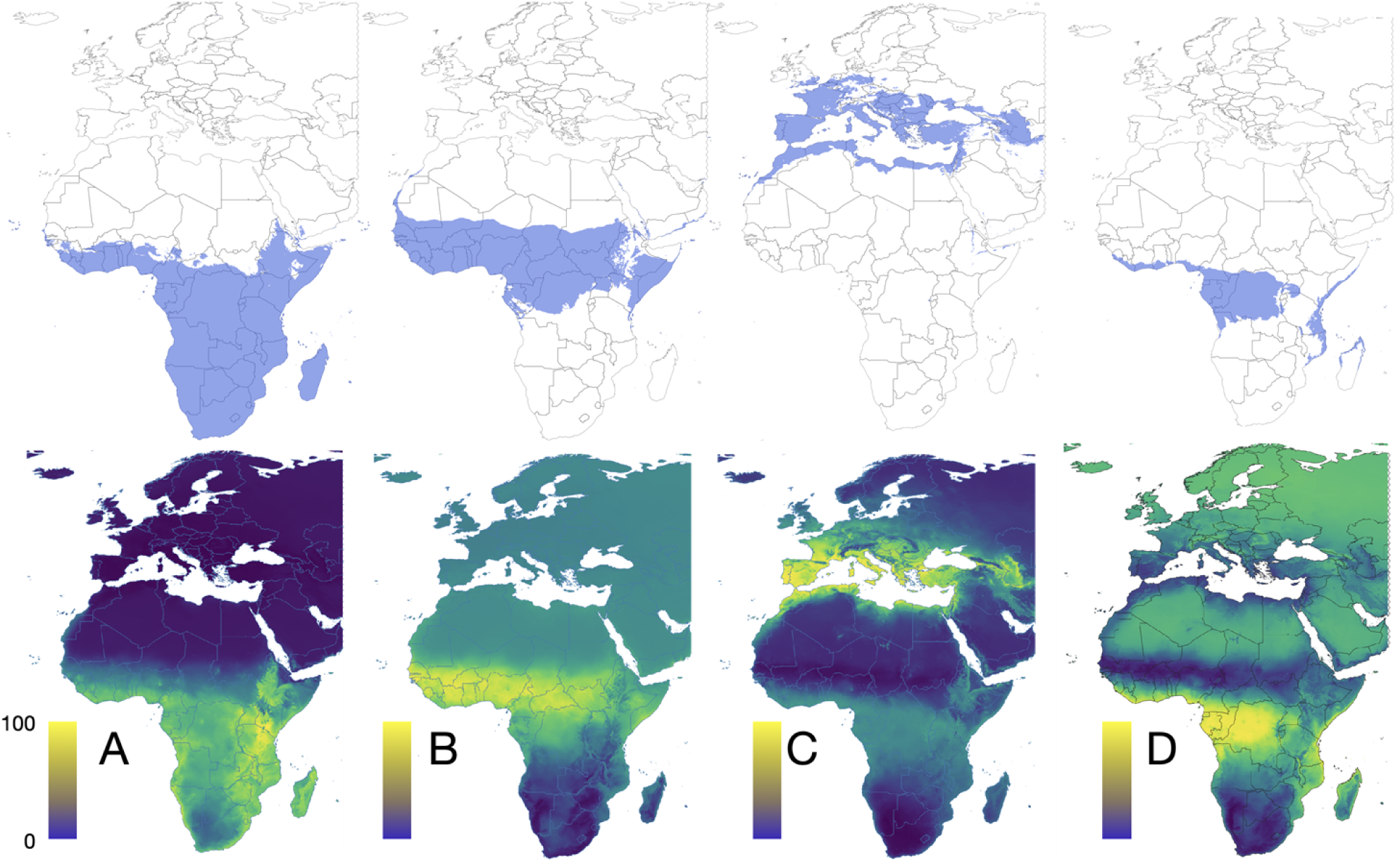
The chorotypes of tick species in the target territory, consecutively numbered #1 to #4 (A to D). Maps in the top row display the range of binomial positive suitability (blue). Maps in bottom row display the probability of occurrence of each chorotype in a range normalized between 0-100. The threshold for binary occurrence has been calculated from the sensitivity and specificity of the predictive mapping of ticks in each chorotype. Supplementary Figure 1 includes the loadings of each tick species in each chorotype.

**Figure 3.**
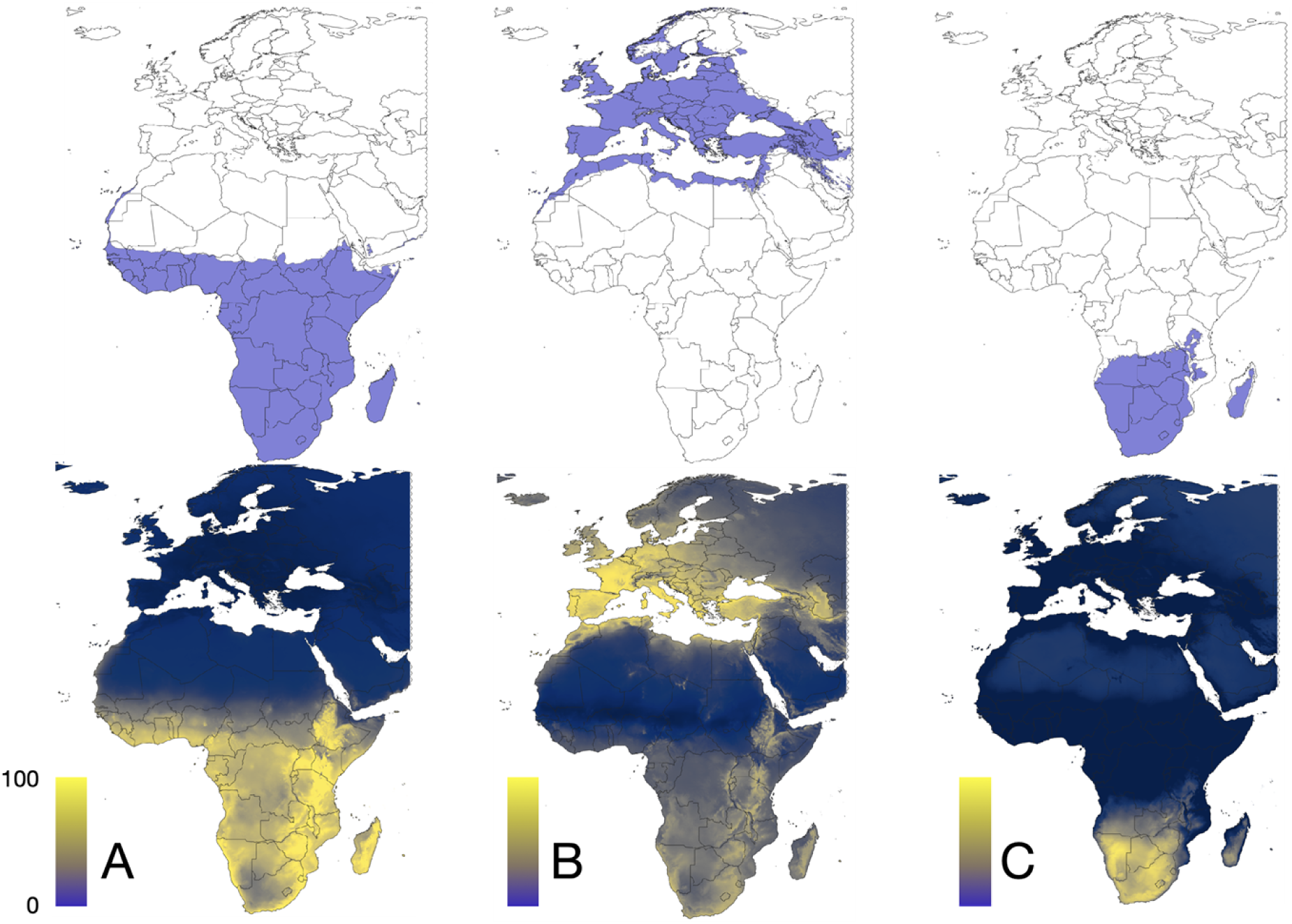
The chorotypes of host genera in the target territory, consecutively numbered #1 to #3 (A to C). Maps in the top row display the range of binomial positive suitability (blue). Maps in bottom row display the probability of occurrence of each chorotype in a range normalized between 0-100. The threshold for binary occurrence has been calculated from the sensitivity and specificity of the predictive mapping of ticks in each chorotype. Supplementary Figure 2 includes the loadings of each host species in each chorotype.

### A climate niche for CCHFV could not be detected

The Non-metric Multidimensional Scaling (NMDS) analysis of 725 points of CCHFV presence and 1,217 pseudo-absences revealed five clusters based on climate covariates across the study region (Figure 4). NMDS failed to discriminate between presence and pseudo-absence points, as all clusters before contained both categories in variable proportions. Cluster 4 contained the highest proportion of positive records (68.3%) in the entire dataset, but these were not correctly separated from pseudo-absence points. The lack of clear separation between presence and pseudo-absence records strongly suggests the absence of a climate niche for CCHFV. Instead, the known range of CCHFV appears to exhibit a patchy distribution that cannot be adequately explained by climate variables alone.

**Figure 4.**
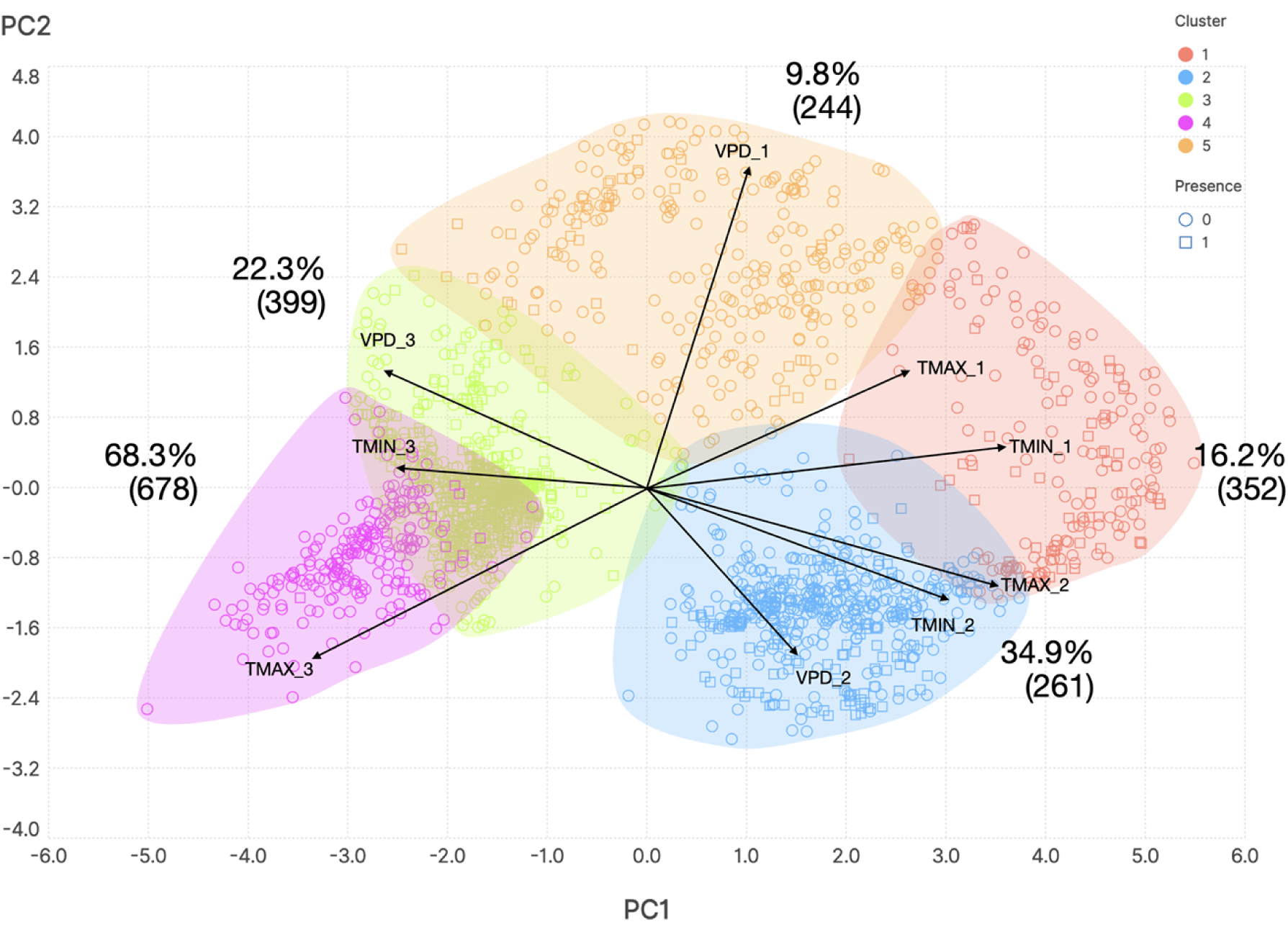
The plot of a non-metric multidimensional scaling (NMDS) in the reduced space, showing 725 records of CCHFV presence and 1,217 pseudo-absences. The NMDS was trained with climate variables to discern a hypothetical climate niche of CCHFV. The optimized output consists of five clusters that explain the climate variability. Positive and negative records of CCHFV could not be separated by the method.

To further explore this, we projected the distribution of CCHFV using only climate variables. The best-performing ensemble (Figure 5) predicted high probabilities of occurrence in the Mediterranean and South Africa. However, the threshold values were unacceptably low, ranging from 0.22 (SVM) to 0.45 (MARS) and 0.44 (GAM).

**Figure 5.**
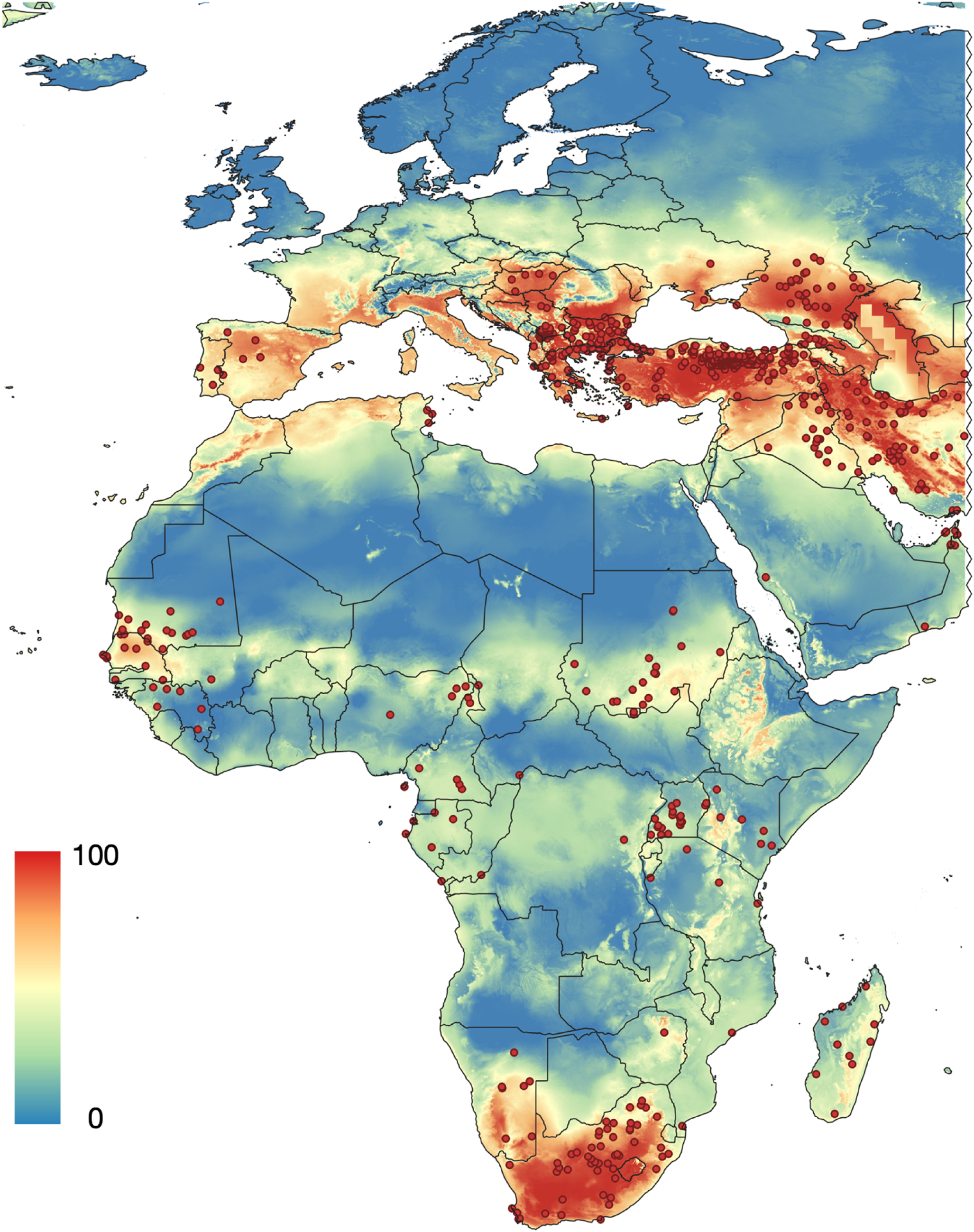
The probability of occurrence of CCHFV calculated only with the climate variables, calculated with a stacking species distribution model (SSDM) based on the best model of the tested algorithms.

The combined ensemble model, based on the best algorithm, tended to overpredict. This resulted in areas of apparent high suitability where no records of the virus or positive serology data exist. Consequently, extensive regions of false positives were found in central Europe, where no records of human cases are known to exist. The specificity values of the best model remained below 0.18, with high omission rates of 0.22–0.31. This approach failed to capture the known distribution of CCHFV in large parts of the Western Palearctic and the Tropics. Therefore, climate alone is not suitable for modelling the distribution of CCHFV.

### Modelling CCHFV occurrence through tick interactions

We employed a RF framework to evaluate the relative contributions of different ticks to the potential distribution of the virus. This is a practical solution to the very large computational task of modelling the range of an organism through the combination of occurrence layers of 82 species of ticks, that must be entered in many combinations, until the best model is obtained.

The RF model correlating available records of CCHFV against the range of the ticks produced good performance values, with a TSS = 1 (the 100% of 1,942 presence/absence of the virus were correctly allocated), RMSE = 0.1434 and LogLoss = 0.1063. Other values of performance of the best model were sensitivity = 0.82, specificity = 0.87, accuracy = 0.84, and Cohen’s kappa = 0.67. The model detected 20 species that are positively correlated with the distribution of CCHFV, included in Figure 6A with the importance as explanatory variable for the model. Notably, the list is composed by all the ticks that bite humans in the target territory, including all the *Hyalomma* spp. that bite humans, plus a selection of ticks that may be important parasites of Artiodactyla. Other species, like *A. sparsum* or *H. aegyptium* parasitize also reptiles but are in the list of important explanatory variables. According to the importance ranked by these studies, *Hyalomma* spp. are confirmed as primary vectors of the virus. The importance of one-host ticks like *R. annulatus* in the transmission of the virus to non-human vertebrates is also noticeable.

**Figure 6.**
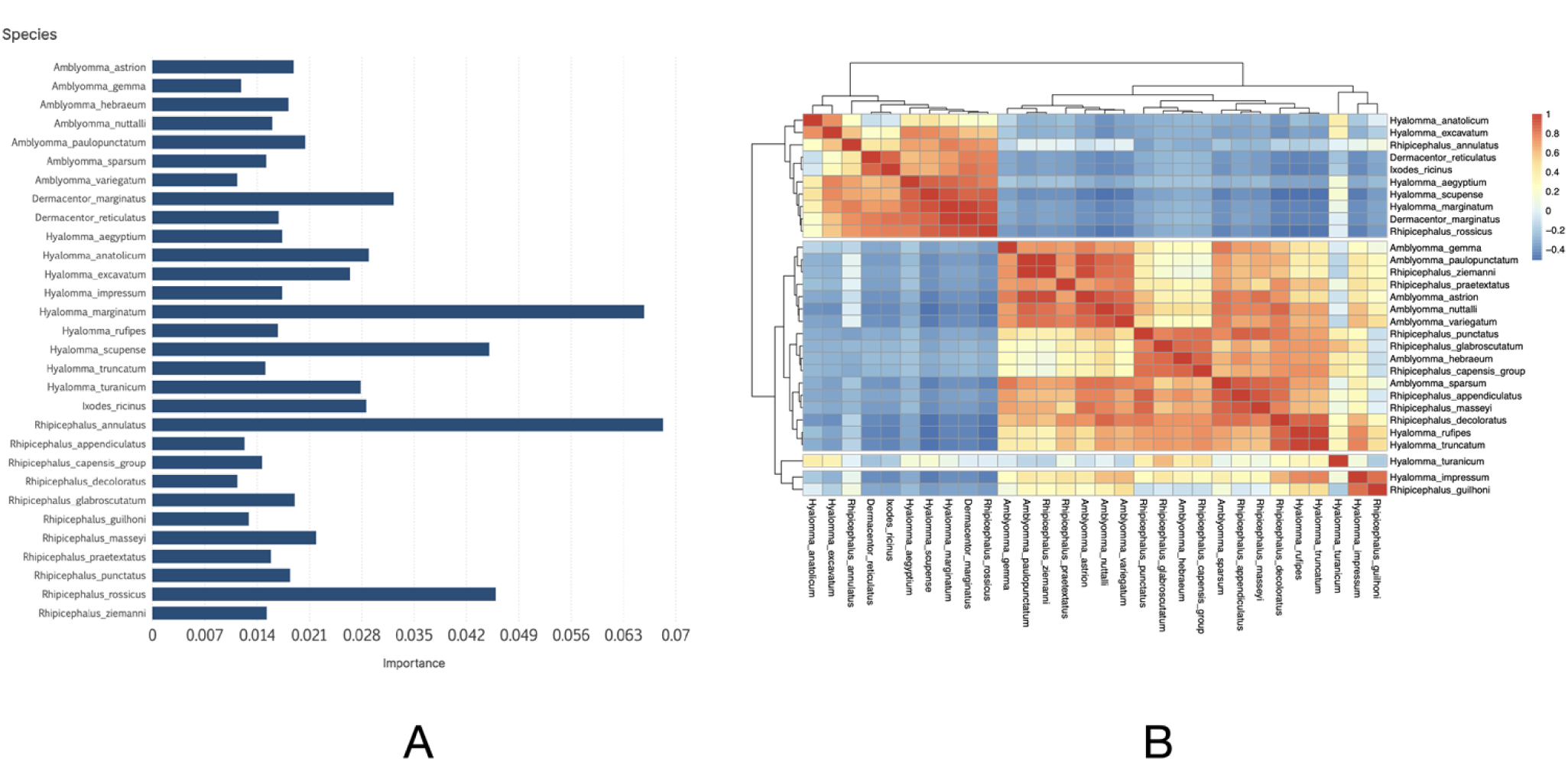
A. The importance of the 30 species of ticks selected as correlated with CCHFV range by an RF model. The species are alphabetically sorted. B. A heatmap displaying the percentage of shared occurrence of the 30 species of ticks in the target territory.

These ticks share their spatial ranges at variable proportions. To understand the correlation of spatial suitability among the tick species, a heatmap was created (Figure 6B). Taxa were entered according to their ordination and grouped by habitat sharing. Since both groups of ticks spread separately in either the Western Palearctic or Afrotropics, the main feature of the heatmap is the almost complete separation of these groups of species into two well defined clusters.

We further projected the RF model using the ranges of the ticks above as explanatory variables to draw the expected range of CCHFV (Figure 7). Being statistically solid as explained before, the RF model detects correctly the known distribution of the positive and negative records. However, the model did not output positive probabilities out of the main clusters of disease occurrence, and the occurrence of CCHFV is projected only for sites where the virus has been already detected.

**Figure 7.**
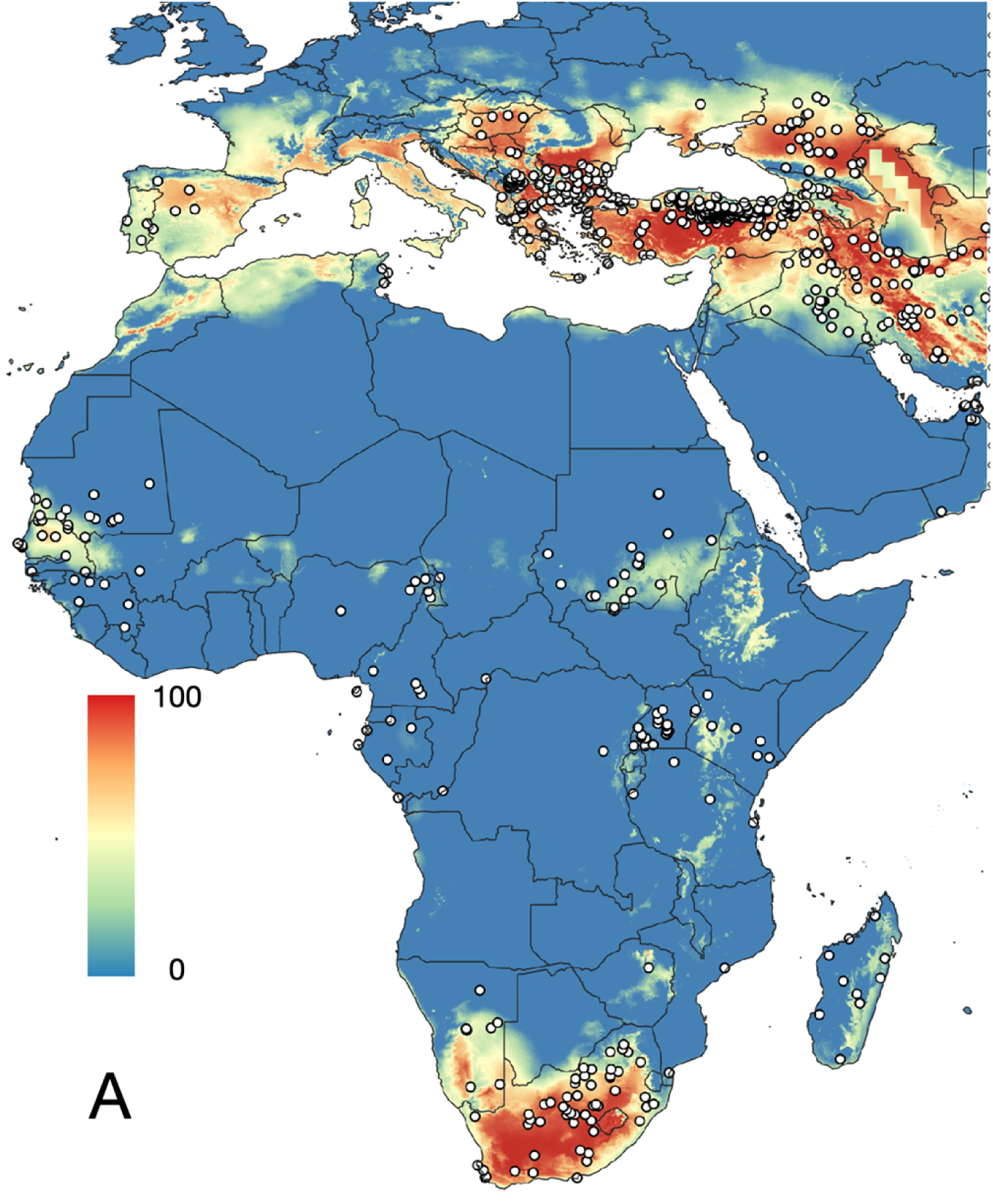
The probability of occurrence of CCHFV using the layers of occurrence of 30 species of ticks as explanatory variables, according to a Random Forest model, trained with the points of CCHFV presence and pseudo-absence and the probability of occurrence of the 82 tick species. The map is projected with the 20 species of ticks detected as correlated with the distribution of CCHFV.

### Explaining CCHFV occurrence through host and tick interactions

We tested more complex models integrating additional ecological layers. Figure 8 presents three approaches: tick chorotypes (Figure 8A), vertebrate chorotypes (Figure 8B), and a combined framework incorporating human-biting ticks, vertebrate chorotypes, livestock density (as a proxy for human–livestock contact), and basic climate covariates (Figure 8C). Model performance metrics are summarised in Table 2.

**Figure 8.**
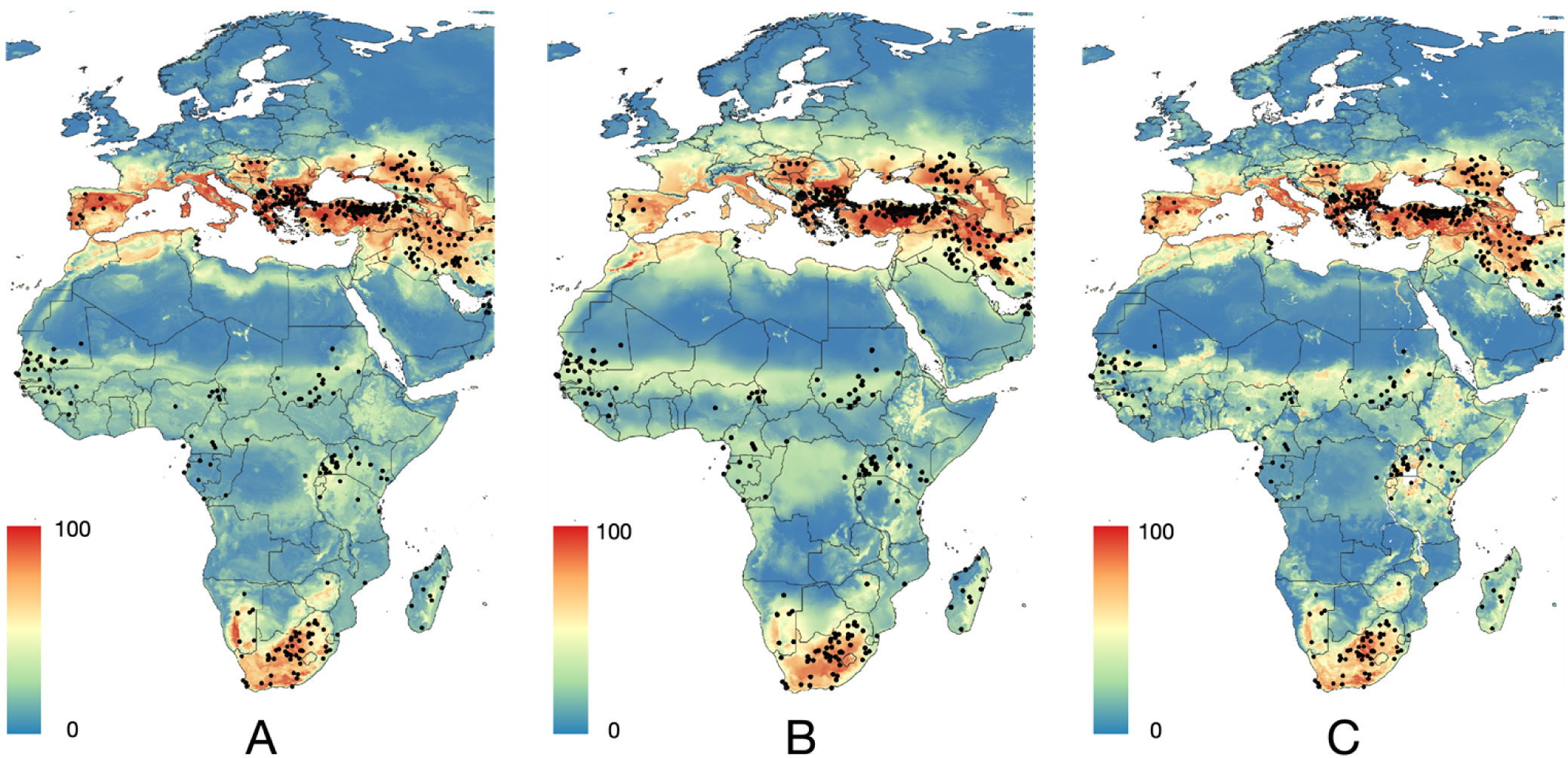
A. Probability of occurrence of CCHFV calculated using the chorotypes of ticks and the climate variables. B. Probability of occurrence of CCHFV calculated with the chorotypes of hosts and the climate variables. C. Probability of occurrence of CCHFV calculated with the distribution of human-biting ticks, the chorotypes of wild hosts and the density of livestock (cattle, sheep, goats). The species of ticks selected are *Amblyomma hebraeum, Amblyomma variegatum, Dermacentor marginatus, Dermacentor reticulatus, Hyalomma aegyptium, Hyalomma anatolicum, Hyalomma excavatum, Hyalomma marginatum, Hyalomma rufipes, Hyalomma scupense, Hyalomma truncatum, Hyalomma turanicum,* and *Ixodes ricinus*.

**Table 2.** Summary of the outcome of algorithms modeling the distribution of CCHFV, with combinations of different explanatory variables. All the combinations included the nine variable species, that were also entered alone, as a final demonstration of the power of climate to discriminate the sites of CCHFV. “HBT+Livestock” refers to the joint use of human-biting ticks, chorotypes of vertebrates, and the density of livestock. “Complete clusters” refers to the simultaneous use of the clusters of ticks and vertebrates. Other categories are self-explained. Threshold indicates the optimal value to convert the habitat suitability map into a binary presence/absence map (higher value is better model). The kappa statistic is a method to evaluate the performance of models predicting presence/absence with a probability.

| Combination of variables and algorithm | Threshold | AUC | Omission rate | Sensitivity | Specificity | Accuracy |
| --- | --- | --- | --- | --- | --- | --- |
| Chorotypes of vertebrates<br>MaxEnt | 0.4383 | 0.8846 | 0.1876 | 0.8123 | 0.8125 | 0.8124 |
| Chorotypes of ticks GAM | 0.4240 | 0.8753 | 0.1903 | 0.8083 | 0.8107 | 0.8097 |
| Chorotypes of ticks MARS | 0.3860 | 0.8803 | 0.1907 | 0.8083 | 0.8100 | 0.8093 |
| Chorotypes of ticks SVM | 0.3740 | 0.8680 | 0.2037 | 0.7954 | 0.7972 | 0.7963 |
| Chorotypes of ticks<br>MaxEnt | 0.3947 | 0.8745 | 0.1949 | 0.8040 | 0.8060 | 0.8051 |
| Only climate<br>GAM | 0.2420 | 0.6619 | 0.3170 | 0.6825 | 0.6833 | 0.6830 |
| Only climate<br>MARS | 0.2010 | 0.6681 | 0.2901 | 0.5917 | 0.6887 | 0.6899 |
| Only climate<br>SVM | 0.3201 | 0.6495 | 0.3057 | 0.6834 | 0.6853 | 0.6843 |
| Only climate MaxEnt | 0.4283 | 0.8598 | 0.2142 | 0.7859 | 0.7858 | 0.7858 |

The best-performing models consistently used either MARS or MaxEnt with the combined explanatory set (Figure 8C), producing the highest sensitivity/specificity balances, lower omission rates, and higher kappa scores. Models using chorotypes alone performed considerably worse, suggesting that the information from chorotypes was insufficient to capture the full complexity of CCHFV ecology and may obscure critical epidemiological relationships.

Overall, our results indicate that the spatial distribution of CCHFV is best explained by a combination of (i) the probability of occurrence of generalist species of ticks, (ii) vertebrate community composition, (iii) livestock density, and (iv) climate covariates. This multidimensional approach outperformed climate-only and tick-only models, reflecting the multifactorial nature of CCHFV transmission and highlighting the importance of integrating host–vector–environment interactions in large-scale epidemiological models.

## Discussion

### 1. Limitations of Climate-Based Models and the Need for Biotic Context

This study presents the most comprehensive analysis to date, integrating abiotic and biotic determinants of the epidemiology of Crimean–Congo haemorrhagic fever virus (CCHFV) to forecast its potential distribution. By compiling reports of human cases, virus isolation sites, and molecular and serological datasets from both humans and animals, we generated and made publicly available a spatially explicit dataset spanning from northern Europe to southern Africa, and from the Atlantic to mid-Asia. Data from central and eastern Russia were intentionally excluded due to inconsistencies in reporting.

Our results challenge the assumption that CCHFV occupies a distinct climatic niche. Models based solely on climatic covariates yielded biased and overly extensive predictions that failed to align with observed occurrences. Climate-only models consistently overestimated northern suitability, a pattern previously noted in studies projecting CCHFV risk in Europe. Such models often identified regions lacking established populations of competent tick vectors and without records of human or animal infection [15, 16, 18, 45]. Consequently, climate-based models alone appear inadequate as proxies for CCHFV epidemiology, as they tend to produce artefactual high-risk areas across central and northern Europe where only sporadic, imported ticks have been detected [46, 47]. To note, the records of the virus have been entered without information about the virus lineage [13] for model training. This is topic worth of research since our results on Multidimensional Scaling showed an unusual number of clusters within the raw records of the virus.

Previous research has largely focused on climate-derived variables as primary determinants of tick distributions and, by extension, CCHFV risk. This emphasis arises from the dual role of ticks as both vectors and reservoirs of the virus [48]. Yet, a wide and poorly characterised range of vertebrate hosts may also act as infection amplifiers [4, 19]. While climate undoubtedly influences tick ecology, recent studies have demonstrated the added importance of landscape and biotic (vertebrate and tick-derived) factors in explaining large-scale CCHFV distribution [14, 49].

Models using only tick distributions as predictors were similarly limited. Although a RF algorithm identified the likely combination of tick species involved in viral circulation, models relying exclusively on tick presence under-predicted the known range of CCHFV. This limitation likely reflects the complexity of co-occurring tick and vertebrate interactions, probably together with the genetic variability of the virus, which obscure individual species’ predictive value. It remains uncertain whether these interactions, or the absence of climatic information, contributed most to model instability. Future modelling efforts should explore these relationships using simplified and interpretable frameworks.

### 2. An Ecological Generalist System Driven by Host–Vector Interactions

The most robust predictions were obtained when integrating distributions of human-biting ticks, livestock density, and host chorotypes. This combination accurately captured approximately 91% of known CCHFV occurrences. Vertebrate chorotypes—clusters of co-occurring species based on range similarity [37, 39] enhanced predictive performance by identifying host assemblages most likely to sustain viral amplification. Notably, no single chorotype is uniquely associated with the virus; rather, each chorotype’s range aligns spatially with sectors of the known CCHFV distribution. These biotic predictors reduced model dimensionality (Reulen & Kneib, 2016) while improving the delineation of both northern and African range limits. Vertebrates likely maintain viral circulation by supporting high tick densities and prolonged viraemia in medium- to large-sized hosts, while immature ticks feeding on small mammals may facilitate local persistence [50]. Incorporating vertebrate community structure therefore bridges vector ecology and pathogen transmission, yielding more ecologically realistic forecasts.

Our findings support an ecological generalist model of CCHFV transmission, as proposed [6]. Under this framework, the virus is not restricted to a narrow set of tick species. Instead, primary human-biting vectors sustain transmission to humans throughout most endemic regions, while secondary tick species maintain circulation among amplifying hosts, predominantly Artiodactyla [27]. These ticks typically rely on Rodentia, Lagomorpha and Insectivora as maintenance hosts for immature stages [20].

Taken together, the presence of CCHFV appears to be a matter of density: more competent tick vectors, more vertebrates capable of sustaining immature ticks, and more hosts that can amplify infection. Human transmission would thus arise from close contact with tick-infested habitats. This system explains the broad geographical extent of human cases without invoking strict molecular coevolution between ticks and vertebrates—an assumption not yet empirically tested.

A variable set of species is likely responsible for CCHFV transmission across different regions. *Hyalomma* spp. consistently emerge as key vectors, together with several *Rhipicephalus* spp. In Europe, transmission is primarily associated with *H. marginatum*, *H. scupense*, and *H. aegyptium*. The role of *D. marginatus* remains unclear, given its partial range overlap with *H. marginatum*; there are no *bona fide* records of CCHFV in *D. marginatus*, as detections refer only to feeding specimens [51]. Our study also provides indirect evidence regarding the potential involvement of *I. ricinus*, long listed among candidate vectors [6]. The northern fringe of predicted suitability coincides with overlap between *I. ricinus* and *H. marginatum*, the latter being the principal human-biting vector in the Mediterranean region. We hypothesise that *I. ricinus* may act as a vector on rare occasions where ranges overlap, *I. ricinus* being a secondary vector unable to sustain the infection in the absence of primary tic vectors like *Hyalomma* spp., although this remains unconfirmed. Laboratory evidence of viral replication in *I. ricinus* cell lines [52] and detections in questing ticks in Turkey [53] warrant further experimental validation.

In central Africa and the Sahel, the combination of human-biting tick presence and livestock density delineated potential transmission hotspots, possibly linked to peri-domestic slaughter or bushmeat practices that increase human exposure. Our approach also identified plausible risk zones in under-studied regions such as Madagascar, where the distribution of *A. variegatum*, the island’s only human-biting tick, matches reports of CCHFV circulation [54]. Overall, our results indicate a widespread viral range, efficiently maintained by ticks feeding on Artiodactyla.

Our best model also highlighted areas in central Africa that may be unsuitable for CCHFV, where ruminant-associated ticks do not overlap with human-biting species, supporting the observed absence of human cases or serological evidence. A further point of uncertainty concerns species such as *R. rossicus* and *R. decoloratus*, which emerged as positively associated with human transmission. The virus has been detected in both species [51], both bite humans [27] and both inhabit regions where human cases occur. At least *R. decoloratus* relies heavily on cattle, suggesting potential for substantial viral circulation. However, these species also overlap with well-established vectors such as *Hyalomma* spp., underscoring the need to clarify their vectorial competence.

The hypothesis of generalist ecological associations contrasts with evidence suggesting potential molecular links between viral lineages and specific tick taxa [1, 55]. Our findings indicate spatial independence between the ranges of major tick vectors in the Western Palaearctic and the Afrotropics. If CCHFV lineages were tightly adapted to particular tick species, cross-regional spread would require genetic adaptation. However, reports of viral strains emerging in unexpected regions support our modelling results and reinforce the view of CCHFV as an ecologically flexible system, potentially facilitated by animal trade and migratory birds [56, 57].

Despite its scope, this study has limitations. Data scarcity across central and eastern Asia hindered comprehensive modelling at the scale of the entire Palaearctic. Furthermore, the spatial resolution of serological data, often aggregated to administrative units, may have introduced minor inaccuracies. Large-scale datasets from central Asia and Africa [58, 59] were excluded due to coarse reporting that precluded consistent spatial matching. Nonetheless, the integration of human and animal serology, human-biting tick distributions, and vertebrate chorotypes produced robust and ecologically coherent predictions, emphasising the value of combining biotic and abiotic dimensions in future epidemiological frameworks.

## Conclusions

Accurate mapping of CCHFV distribution requires a multifactorial approach that integrates both abiotic and biotic predictors. Models based solely on climate or on the ranges of known tick vectors generate biased outputs. Our findings indicate that CCHFV lacks a strict climatic niche and that its persistence depends on the co-occurrence of vector and vertebrate communities. Incorporating human-biting tick distributions, livestock density, and vertebrate chorotypes significantly improved predictive performance, providing a mechanistic understanding of the ecological processes sustaining virus circulation. This integrative framework establishes a robust foundation for forecasting CCHFV risk and highlights the importance of combining high-resolution ecological, vector, and host data with environmental variables in future studies of vector-borne zoonoses.

## Supporting information

**Supplementary Table 1**: **The species of ticks included in this study**. Notes about its importance as human biting ticks have been included using the same terms as in the original compilation (Guglielmone and Robbins, 2018). Parasitism on Artiodactyla is included since medium and large ungulates are considered important vertebrates contributing to amplification of CCHFV. Categories of the prevalence of ticks on these groups of hosts are identical to the original report. Ticks belonging to the genus *Haemaphysalis* were originally included in the study but later removed because the lack of association with the records of CCHFV.

**Supplementary Table 2**. **The compilation of serological surveys from literature**. The file is organized in three sheets of information and is self-explained. It includes one sheet for data compiled from humans, another sheet for data generated from animals, and a third one that includes the rejected papers in the bibliographical query and revision.

**Supplementary Figure 1**: **The representation in the reduced space of the chorotypes of the ticks species included in this study**. Dots are placed along the coordinates of chorotypes 1 and 2, with colour explaining the ordination in the chorotype 3. The chart intends to be informative but not exhaustive as some points are very near in the reduced space and their labels could not be separated.

**Supplementary Figure 2**: **The representation in the reduced space of the chorotypes of the genera of vertebrates included in this study**. Dots are placed along the coordinates of chorotypes 1 and 2, with colour explaining the ordination in the chorotype 3. The chart intends to be informative but not exhaustive as some points are near in the reduced space and their labels could not be separated.

